# Reduced Hippocampal and Amygdala Volumes as Mechanisms of Stress Sensitization to Depression following Childhood Violence Exposure

**DOI:** 10.1101/798033

**Authors:** David G. Weissman, Hilary K. Lambert, Alexandra M. Rodman, Matthew Peverill, Margaret A. Sheridan, Katie A. McLaughlin

## Abstract

**OBJECTIVE:** Stressful life events are more likely to trigger depression among individuals exposed to childhood adversity. However, the mechanisms underlying this stress sensitization remain largely unknown. Any such mechanism must be altered by childhood adversity and interact with recent stressful life events, magnifying their association with depression. This study investigated whether reduced hippocampal and amygdala volumes are mechanisms of stress sensitization following childhood violence exposure.

**METHODS:** A sample of 149 youth (aged 8-17 years), with (N=76) and without (N=73) exposure to physical abuse, sexual abuse, or domestic violence participated. Participants completed a structural MRI scan and were assessed for symptoms of depression. Approximately two years later, stressful life events were assessed along with depression symptoms in 120 participants (57 violence-exposed).

**RESULTS:** Childhood violence exposure was associated with smaller hippocampal and amygdala volumes. Stressful life events occurring during the follow-up period predicted worsening depression over time, and this association was magnified among those with smaller hippocampal and amygdala volumes. Significant moderated mediation models revealed indirect effects of violence exposure on increasing depression over time through hippocampal and amygdala volumes, particularly among youths who experienced more stressful life events.

**CONCLUSIONS:** These results provide novel evidence for reduced hippocampal and amygdala volumes as mechanisms of stress sensitization to depression following exposure to violence. These findings suggest that hippocampal and amygdala-mediated emotional and cognitive processes may contribute to vulnerability to stressful life events following childhood violence exposure.

## Introduction

Childhood adversity is common, and is strongly associated with psychopathology, including depression (1,2). Heightened sensitivity to stress has been identified as a pathway through which childhood adversity may increase risk for depression, such that stressful life events are more likely to trigger depressive episodes in those who have experienced childhood adversity (3). This process—called stress sensitization—has been observed in adults (1) and adolescents (4), and in both population-based (1) and longitudinal studies (3,4). Despite consistent evidence for stress sensitization to depression following childhood adversity, the mechanisms that underlie this process have rarely been investigated and remain largely unknown. Understanding the psychological and biological mechanisms of stress sensitization may inform targets for interventions to prevent and treat depression and reduce stress vulnerability in individuals exposed to childhood adversity.

In order to qualify as a mechanism of stress sensitization, a specific psychological or neurobiological marker must: 1) be associated with childhood adversity, and 2) interact with recent stressful life events, strengthening their association with depression. We propose that alterations in hippocampal and amygdala volumes are potential mechanisms of stress sensitization. In a recent systematic review, we found that smaller amygdala volume and smaller hippocampal volume are the most consistent differences in brain structure associated with threat-related adversity in children and adolescents (5). Threatening experiences early in life (e.g., interpersonal violence) may alter amygdala structure and function to promote rapid identification of threats in the environment (6). Changes in emotional information processing may be adaptive in dangerous environments, but maladaptive in safe environments or in the face of less severe stressors. Smaller amygdala and hippocampal volumes are associated with increased physiological reactivity to stress and disrupted feedback of physiological stress responses respectively (7,8). Further, increased reactivity to threat and difficulties with emotion regulation have been proposed as potential psychological mechanisms of stress sensitization (9).

Smaller hippocampal volume is associated with depression in adults in meta-analysis (10). Evidence of a direct relation between hippocampal volume and depression is less consistent in children and adolescents, with several studies observing a negative association and others finding no association (11,12), or even a positive association (13). Likewise, relations between amygdala volume and depression are mixed in children and adolescents (11,12,14). These discrepant patterns are consistent with the possibility that the association between hippocampal and amygdala volumes and depression are moderated by factors such as age or stressful life events. In the current study, we examine whether hippocampal and amygdala volumes moderate the association of stressful life events with depression, such that stressors are more likely to predict increases in depression over time among youth exposed to violence and with smaller hippocampal and amygdala volumes.

To our knowledge, little or no research to date has investigated the role that reduced amygdala volume plays in magnifying vulnerability to depression following stressful life events among those with a history of violence exposure. There is some evidence, however, that reduced hippocampal volume may increase vulnerability to stressful life events. Maternal aggression leads to greater increases in depressive symptoms over time among early adolescents with smaller hippocampal volume (15). In addition, one study found that young adults exposed to violence in childhood had smaller hippocampal volume, which interacted with recent stress in predicting anxiety, such that recent stressors were only associated with elevated anxiety among participants with small hippocampal volume (16). This evidence is promising in suggesting that reduced hippocampal volume may be a mechanism of stress sensitization to depression in children and adolescents with a history of violence exposure, although this possibility is largely untested.

The current study utilizes a prospective longitudinal design to evaluate whether reduced hippocampal and amygdala volumes are mechanisms underlying stress sensitization to depression in children and adolescents exposed to violence. We expected that the associations of stressful life events with increases in depression symptoms over time would be stronger among participants exposed to violence, indicating stress sensitization. In addition, we predicted that youth exposed to violence would have smaller hippocampal and amygdala volumes than those who had never been exposed to violence. We further hypothesized that hippocampal and amygdala volumes would interact with stressful life events in predicting depression both concurrently and prospectively, such that stressful life events would be more strongly associated with depression symptoms among participants with smaller hippocampal and amygdala volumes. Finally, we estimated moderated mediation models that evaluated whether an indirect effect of violence exposure on depression symptoms through hippocampal and amygdala volumes, was magnified in the context of stressful life events.

## Method

### Participants

Participants were 160 children and adolescents between the ages of 8 and 17 living in the Seattle area who were recruited for a study examining neural development in youths with and without exposure to violence. Youth and caregivers were recruited for participation at schools, after-school and prevention programs, adoption programs, food banks, shelters, parenting programs, medical clinics, and the general community in Seattle, WA between January 2015 and June 2017. Recruitment efforts were targeted at recruiting a sample with variation in exposure to violence. To do so, we recruited from neighborhoods with high levels of violent crime, clinics that served a predominantly low-SES catchment area, and agencies that work with families who have been victims of violence (e.g., domestic violence shelters, programs for parents mandated to receive intervention by Child Protective Services). Inclusion criteria for the violence-exposed group included exposure to physical or sexual abuse or direct witnessing of domestic violence. Children in the control group were matched to children in the violence-exposed group on age, sex, and handedness; inclusion criteria required an absence of exposure to significant interpersonal violence. Exclusion criteria included IQ < 80, presence of pervasive developmental disorder, active psychotic symptoms or mania, active substance abuse, and presence of safety concerns. Participants who completed the MRI visit also met standard MRI inclusion criteria (i.e., absence of braces, claustrophobia). Written informed consent was obtained from legal guardians while children provided written assent.

Eleven participants were excluded from further analysis due to excessive head movement, resulting in 149 participants (76 violence-exposed) with usable data on hippocampal volume. A total of 120 (57 violence-exposed) of the 149 participants with usable data returned for follow-up assessments of stressful life events and depression approximately two years later (*M* = 643 days, *SD =* 222, 81% retention rate). The 29 participants for whom follow-up data was missing did not differ significantly in the proportion who were violence exposed (18 violence exposed, 11 unexposed, *χ*^2^ = 1.44, *p* = .230), hippocampal (*d* = .060, *t* = 0.30, *p* = .766) or amygdala volume (*d* = .045, *t* = 0.21, *p* = .832), baseline depression symptoms (*d* = .299, *t* = 1.15, *p* = .249), or stressful life events at baseline (*d* = .352, *t* = 1.56, *p* = .127). See Table 1 for socio-demographic characteristics of the final sample.

**Table 1:**
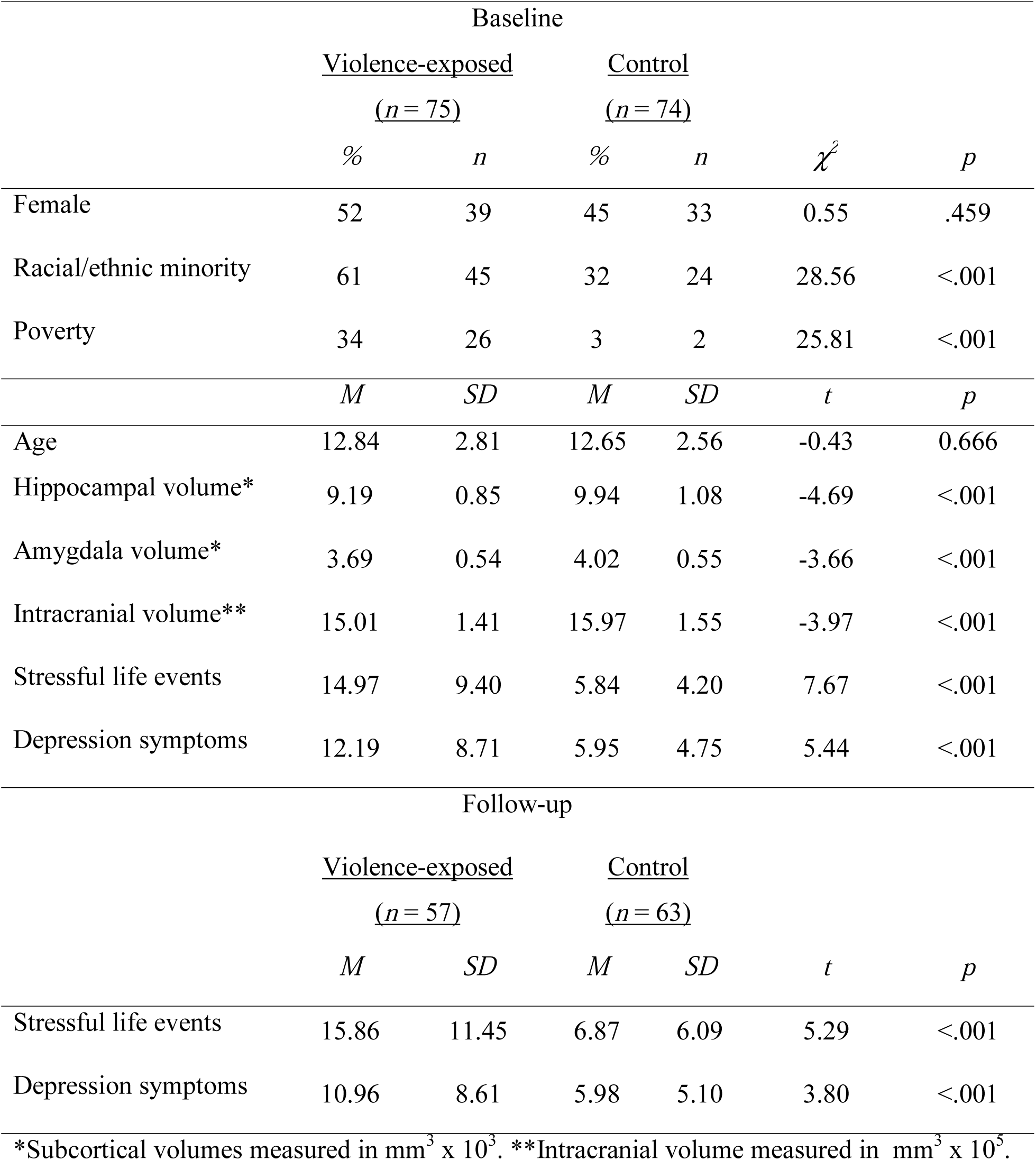
Distribution of study variables by violence exposure.

### Measures

#### Violence Exposure

A multi-informant, multi-method approach was used for assessing exposure to violence. Youths completed the Childhood Experiences of Care and Abuse (CECA) interview, which assesses caregiving experiences, including physical and sexual abuse (17). We modified the interview to ask parallel questions about witnessing domestic violence (i.e., directly observing violence directed at a caregiver). The Childhood Trauma Questionnaire (CTQ) is a 28-item self-report scale that assesses the frequency of maltreatment during childhood, including physical and sexual abuse (18) and was also completed by children. The UCLA PTSD Reaction Index (PTSD-RI) was completed by both the child and a caregiver. It includes a trauma screen that assesses exposure to numerous traumatic events, including physical abuse, sexual abuse, and domestic violence and additionally assesses PTSD symptoms (19). The Juvenile Victimization Questionnaire (JVQ) was completed by caregivers, and includes 34 items assessing exposure to crime, child maltreatment, peer and sibling victimization, sexual victimization, and witnessing and indirect violence (20). Children were classified as experiencing physical or sexual abuse if abuse was endorsed by the child on the CECA, PTSD-RI trauma screen, or above the validated threshold on the CTQ (21) or reported by the parent on the JVQ or PTSD-RI. Domestic violence was assessed by child-report only on the CECA interview and PTSD-RI.

#### Depression

Symptoms of depression were assessed at both baseline and follow-up with the Children’s Depression Inventory-2 (CDI-2), a recently revised version of the widely used self-report measure of depressive symptoms in children and adolescents (22). The CDI-2 is a 27-item scale. For each item, participants choose between 3 statements that correspond to a 3-point scale. Higher scores indicate more severe depression. The CDI-2 demonstrated excellent internal consistency in our sample (α = .89).

#### Stressful life events

Stressful life events occurring in the past year were assessed at both baseline and follow-up using the UCLA Life Stress Interview (23), a semi-structured interview designed to characterize stressful life events as objectively as possible. The interview has been extensively validated, adapted for use with children and adolescents, and is widely considered to be the gold standard approach for assessing stressful life events (3,24).

The interview uses a series of structured prompts to query numerous domains of the child’s life (peers, parents, household/extended family, neighborhood, school, academic, health, finance, and discrimination). An independent research team objectively codes the likely impact of each event for a child of that age and sex from 1 (no negative impact) to 5 (extremely severe negative impact) scale, including half-points (e.g., 1.5, 2.5). These objectively coded impact scores reduce concerns about recall bias or differences in perceptions of stress that can artificially inflate associations between stressful life events and psychopathology. Child participants and a parent independently completed the interview and reported on stressful life events experienced by the child in the past year. Following prior work, a total impact score was computed by taking the sum of the impact scores of all reported events (excluding those coded as 1), which provides a weighted average of both the number and severity of stressors that occurred (3). Higher scores indicate a greater frequency and/or severity of stressful life events in the past year.

The interview was administered at both the baseline and follow-up visits, each time probing stressful life events occurring in the prior year. Total impact scores based on events reported by parents and children were moderately correlated at both baseline (*r* = .56) and follow up (*r* = .49). The higher of the child and parent total impact score was used. At baseline, child reported scores were used for 44 participants, parent reported scores were used for 88 participants, and parent and child reports were the same for 17 participants. At follow-up, child reported scores were used for 36 participants, parent reported scores were used for 69 participants, and parent and child reports were the same for 15 participants.

#### Poverty

Parent-reported family income was compared to the poverty threshold for a family of that size as indicated by the U. S. Census Bureau (25), to determine if families were above or below the poverty threshold. Family income was not reported for 10 participants (8 violence-exposed). For missing cases, poverty was imputed by multiple imputation (100) using the *mice* package in R (26).

### Structural MRI acquisition

Scanning was performed on a 3T Phillips Achieva scanner at the University of Washington Integrated Brain Imaging Center using a 32-channel head coil. T1-weighted MPRAGE volumes were acquired (repetition time = 2530 ms, TE=3.5ms, flip angle=7°, FOV=256×256, 176 slices, in-plane voxel size=1mm^3^).

### Structural MRI processing

Measures of left and right hippocampal and amygdala volumes and total intracranial volume were obtained using automatic segmentation in FreeSurfer 5.3. Each segmentation was inspected manually for over or under inclusion of tissue in the labelled cortex by at least two investigators. Eleven participants were excluded from further analysis due to artifacts from excessive head movement which could not be remediated. No manual edits were performed on subcortical segmentations. Given, that there is no consistent pattern of lateralization in findings of the relation between childhood violence exposure and hippocampal or amygdala volume (6,27,28), and to reduce multiple comparisons, right and left volumes were summed to create measures of bilateral hippocampal and bilateral amygdala volume. To keep variables on a similar scale for regression analyses, hippocampal and amygdala volumes were all divided by 1,000, and intracranial volumes were divided by 100,000.

### Data Analysis

Linear regressions were conducted in R to test for stress sensitization. We evaluated the relation between violence exposure and depression at baseline, controlling for poverty and racial/ethnic minority status. We then evaluated whether a pattern of stress sensitization was present with depression as the outcome variable, stressful life events, violence exposure, and their interaction as predictors, and poverty, racial/ethnic minority status, sex, and age as covariates of non-interest. We repeated these analyses, with depression at follow-up as the outcome variable, stressful life events at follow-up, violence exposure, and their interaction as predictors, and depression at baseline, poverty, racial/ethnic minority status, sex, age, as covariates of non-interest.

To determine whether hippocampal and amygdala volumes are mechanism of stress sensitization, we first evaluated whether violence exposure was associated with each, controlling for intracranial volume, poverty, and racial/ethnic minority status. Next, we examined whether hippocampal and amygdala volumes were associated with depression, controlling for poverty, racial/ethnic minority status, sex, age, and the time elapsed from baseline to follow-up. We then examined the association between hippocampal/amygdala volume, stressful life events at follow-up, and their interaction as predictors of depression at follow-up, controlling for violence exposure, depression at baseline, poverty, racial/ethnic minority status, sex, age at baseline, and the time elapsed from baseline to follow-up. To determine if effects were specific to hippocampus and amygdala, we also conducted control analyses using thalamus volume. Finally, moderated mediation models were tested with bootstrapped confidence intervals (10,000 iterations) using version 2.13 of the process macro in SPSS. Data and analysis code are available at https://github.com/dgweissman/stress-sensitization.

## Results

### Descriptive statistics

Descriptive statistics of study variables and demographic characteristics are summarized for violence-exposed and unexposed youth in Table 1. As expected, violence-exposed and unexposed youth differed on all variables of interest.

### Stress sensitization

The association between stressful life events and depression symptoms was not moderated by violence exposure at baseline (*B* = -.042*, S.E.* = .193*, t* = −0.22*, p* = .828) or follow-up (*B* = .165*, S.E.* = .130*, t* = 1.27*, p* = .208). Although the interaction between violence exposure and stressful life events was not significant in predicting depression symptoms, stratified correlation analyses demonstrate a strong positive zero-order association between stressful life events and depression symptoms at follow-up, among violence-exposed youths but no association among controls (See Figure 1).

**Figure 1:**
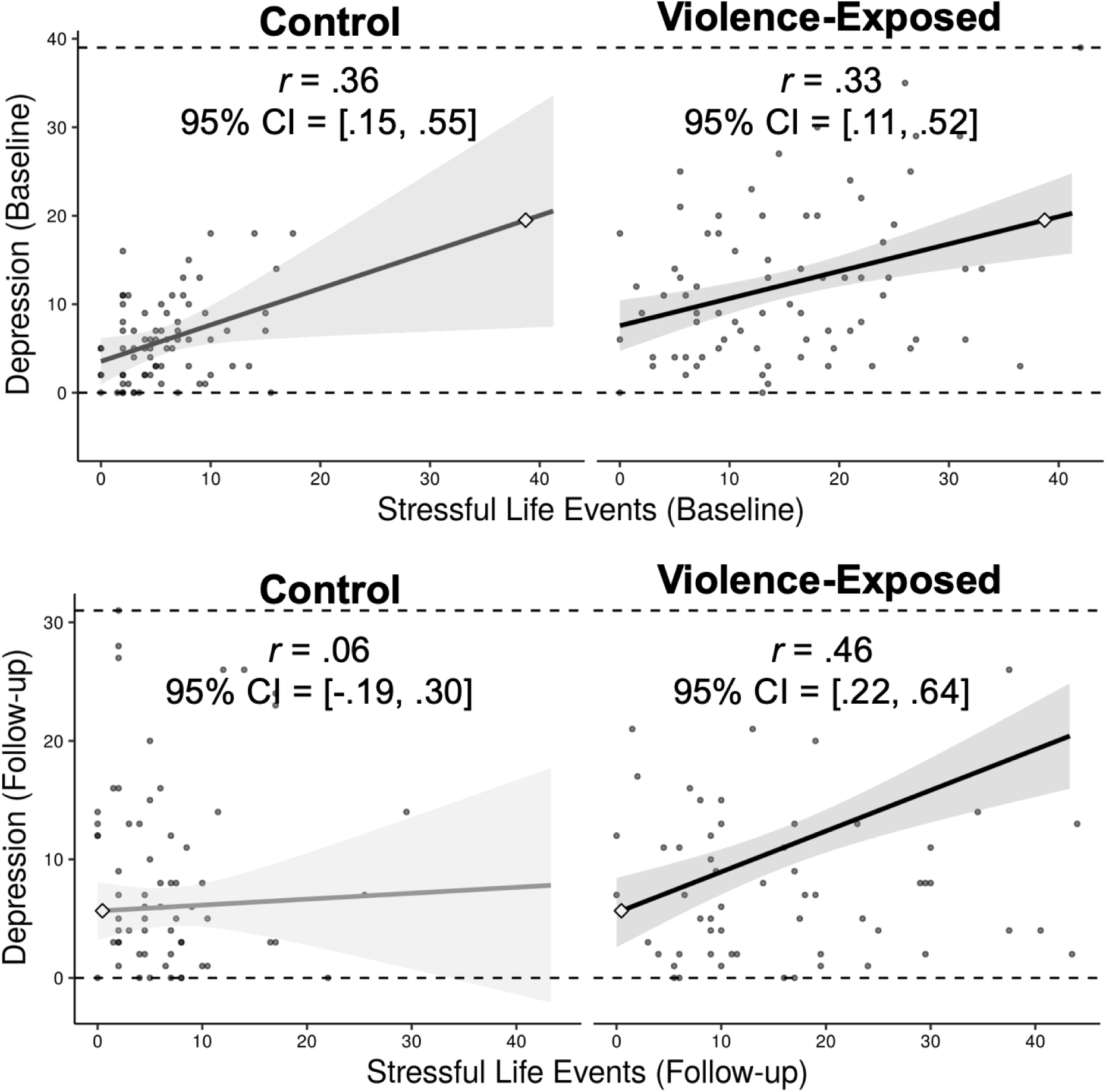
Associations between stressful life events and depression within the violence-exposed and control groups. Stressful life events were significantly correlated with concurrent depression at baseline among violence-exposed and control youth. Stressful life events were significantly correlated with concurrent depression at follow-up among violence-exposed but not control youth.

### Violence exposure and subcortical volumes

Violence exposed participants had significantly smaller hippocampal (*B* = -.364, *S.E.* = .167, *t* = −2.18, *p* = .031) and amygdala (*B* = -.193, *S.E.* = .097, *t* = −2.00, *p* = .048) volumes than control participants. Thalamus volume did not differ between violence exposed and unexposed participants (*B* = -.025, *S.E.* = .179, *t* = −0.14, *p* = .889).

### Subcortical volumes and depression

Hippocampal volume was not associated with depression at baseline (*B* = .587, *S.E.* = .658, *t* = 0.89, *p* = .374), or follow-up (*B* = -.568, *S.E.* = .622, *t* = −0.91, *p* = .364). Amygdala volume was not associated with depression at baseline (*B* = -.983, *S.E.* = 1.144, *t* = −0.86, *p* = .391) or follow-up (*B* = −1.03, *S.E.* = 1.10, *t* = −0.94, *p* = .349).

### Stressful Life Events Interact with Subcortical Volumes

Stressful life events occurring during the follow-up period interacted with hippocampal volume (*B* = -.118, *S.E. =* .057, *t* = −2.06, p = .042) and amygdala volume (*B* = -.222, *S.E.* = .102, *t* = −2.18, p = .032) in predicting depression at follow-up (Table 2). The interaction between thalamus volume and stressful life events in predicting depression was not significant (*B* = -.014, *S.E. =* .044, *t* = −0.32, p = .752). Simple slopes revealed that the association between stressful life events and increases in depression symptoms over time was strongest among participants with small to average hippocampal and amygdala volumes and not significant among participants with larger hippocampal and amygdala volumes (Figure 2).

**Table 2:** Interaction of stressful life events with hippocampal and amygdala volume in relation to depression.

| *Predictors* | **Depression (Follow up)** |  |  |  |  |  |  |  |
| --- | --- | --- | --- | --- | --- | --- | --- | --- |
|  | Hippocampus |  |  |  | Amygdala |  |  |  |
|  | *B* | *SE* | *t* | *p* | *B* | *SE* | *t* | *p* |
| Sex | 1.15 | 1.29 | 0.89 | .375 | 1.16 | 1.29 | 0.90 | .370 |
| Age | .173 | .210 | 0.82 | .412 | .257 | .207 | 1.24 | .217 |
| Time elapsed (years) | -.573 | .840 | -0.68 | .497 | -.615 | .840 | -0.73 | .465 |
| Racial/ethnic minority | 0.15 | 1.12 | 0.14 | .891 | 0.15 | 1.13 | 0.14 | .892 |
| Poverty | -1.39 | 1.49 | -0.93 | .355 | -0.86 | 1.50 | -0.57 | .568 |
| Depression (Baseline) | **.568** | **.080** | **7.05** | **<.001** | **5.33** | **.082** | **6.52** | **<.001** |
| Violence Exposure | 0.57 | 1.42 | 0.59 | .556 | 1.32 | 1.38 | 0.96 | .339 |
| Intracranial Volume | .706 | .512 | 1.38 | .171 | .538 | .491 | 1.10 | .275 |
| Stressful Life Events (SLE) | **.142** | **.060** | **2.37** | **.020** | .092 | .057 | 1.62 | .108 |
| Subcortical Volume (SV) | -1.05 | 0.63 | -1.66 | .099 | -1.39 | 1.08 | -1.28 | .103 |
| SV x SLE | **-.118** | **.057** | **-2.06** | **.042** | **-.222** | **.102** | **-2.18** | **.032** |

**Figure 2:**
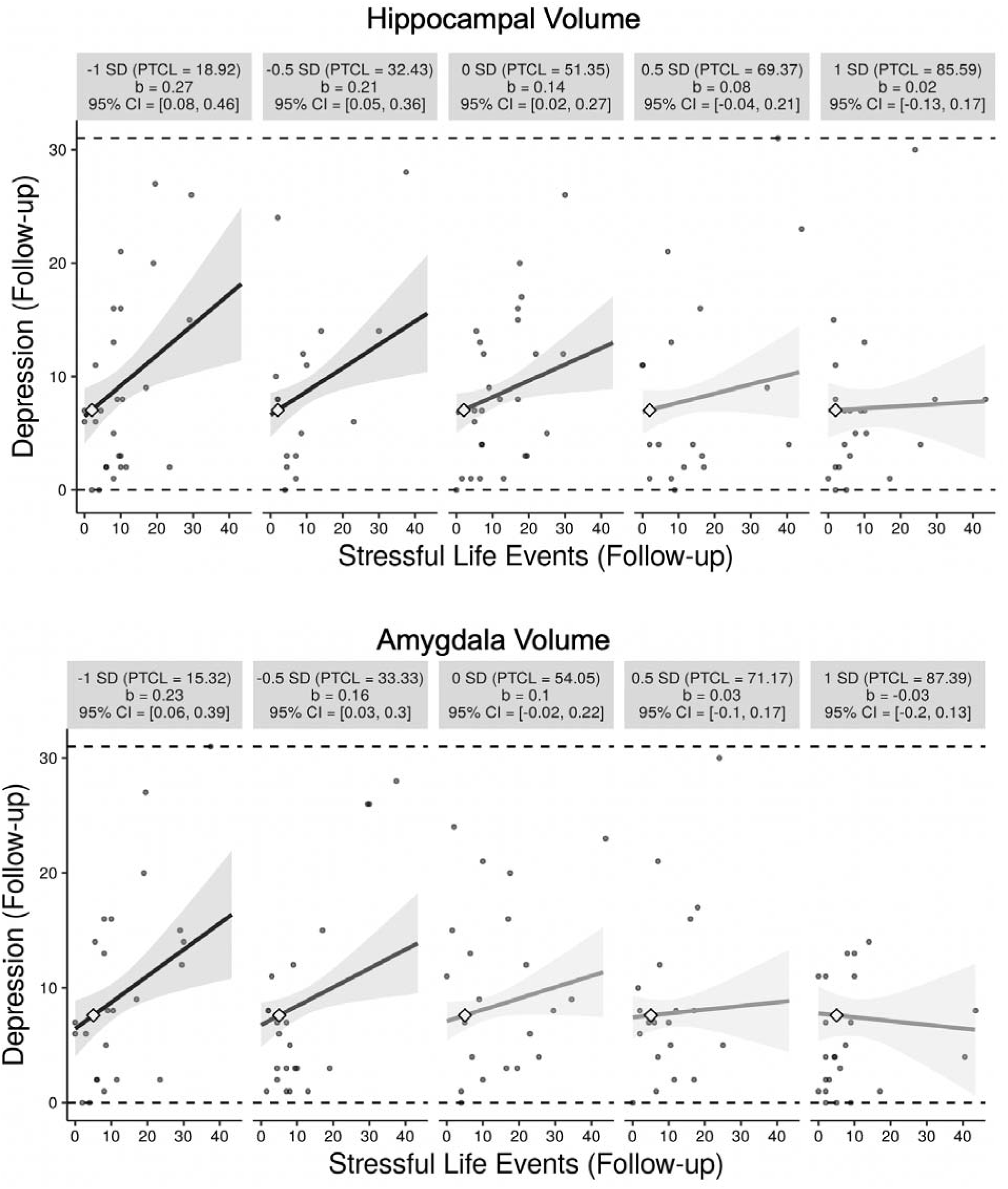
Moderation of the relation between stressful life events and depression by hippocampal and amygdala volume. Stressful life events are positively associated with depression symptoms at follow-up, controlling for baseline depression, in participants with small to average hippocampal volume but not larger (.5 SD above the mean and higher) hippocampal volume and in participants with small (-.5 SD below the mean and smaller) but not average or larger amygdala volume. Figure produced using the interActive data visualization tool (35).

### Moderated Mediation

Moderated mediation analyses revealed a conditional indirect effect of violence exposure on depression at follow up, via hippocampal volume, conditional on recent stressful life events (Figure 3). Analyses also revealed a conditional indirect effect of violence exposure on depression at follow up, via amygdala volume, conditional on recent stressful life events (Figure 3). The bootstrapped 95% confidence interval of the standardized index of moderated mediation for the model with hippocampal volume (.061, 95% CI .010 to .149) and the model with amygdala volume (.064, 95% CI .011 to .160) did not include zero, suggesting a significant conditional indirect effect.

**Figure 3:**
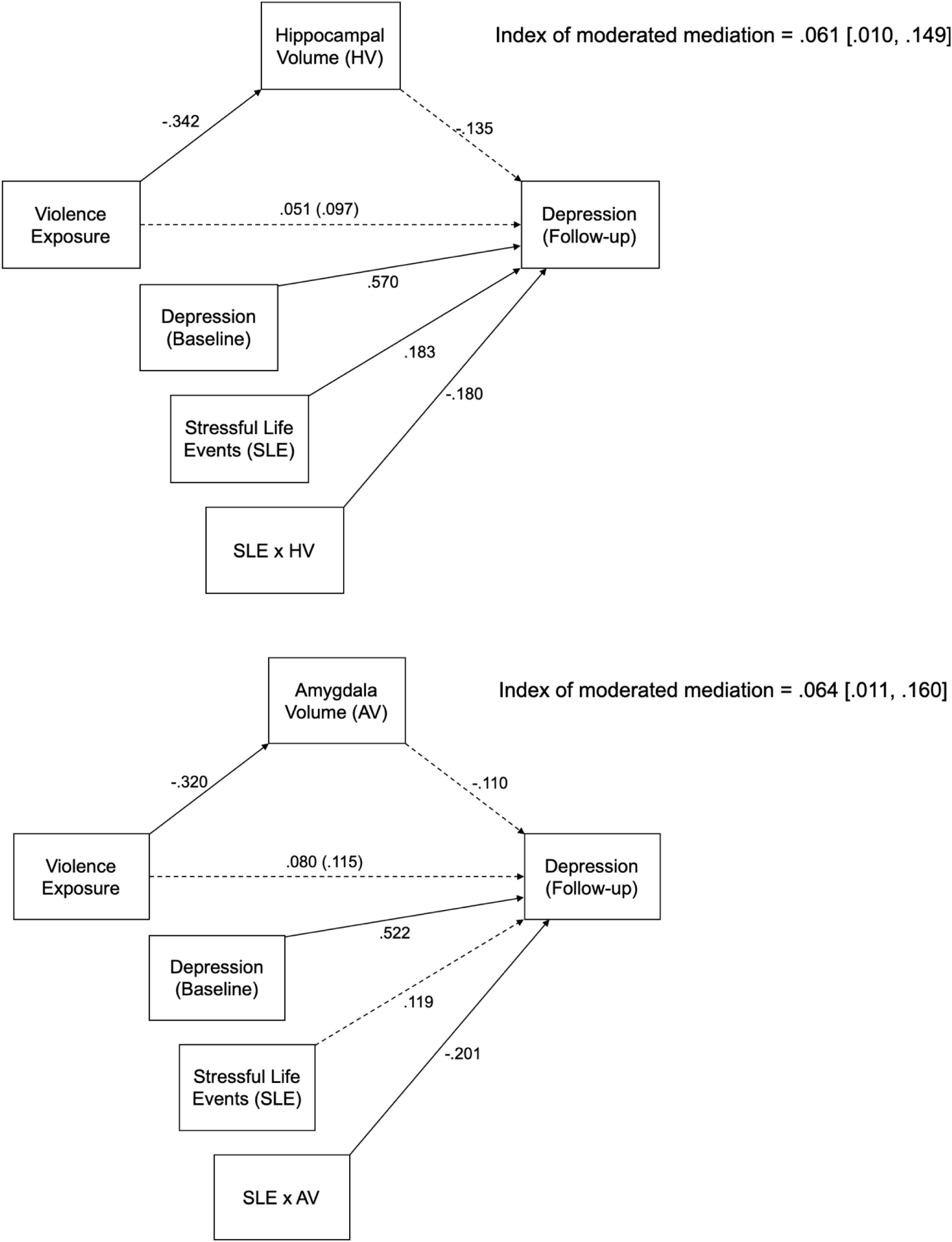
Moderated Mediation Model Results. Results of regression-based moderated mediation models of the indirect effect of violence exposure on follow-up depression via hippocampal and amygdala volume, conditional on recent stress, and controlling for baseline depression. Sex, age, time elapsed between visits, poverty, racial/ethnic minority status, and total intracranial volume are not shown, but were also included as covariates. All coefficients are standardized. 95% bootstrapped confidence intervals are based on 10,000 iterations. Solid lines indicate significant relations. Dotted lines indicate nonsignificant relations.

## Discussion

Children and adolescents who have experienced adversity, including violence exposure, are at elevated risk for depression after stressful life events later in life (4,29). This study provides evidence that smaller hippocampal and amygdala volumes among youths exposed to violence are potential mechanisms underlying this pattern of stress sensitization. We replicate prior work demonstrating that violence-exposed children and adolescents have smaller hippocampal and amygdala volumes and extend it by demonstrating that these reductions in hippocampal and amygdala volumes magnify the association of stressful life events with the emergence of depressive symptoms in a longitudinal design. This provides compelling evidence that differences in hippocampal and amygdala structure following violence exposure may alter the way youth respond to stressors in their environment, contributing to risk for depression.

Although the association between recent stressful life events and increases in depression symptoms over time was significant and positive among violence exposed youth and absent among control participants, violence exposure did not interact with stressful life events to predict increases in depression symptoms over time. However, this pattern is well-replicated in larger samples (1,3,4) and broadly consistent with the pattern of associations between stressful life events and depression symptoms observed here.

Our findings are consistent with prior studies examining the relation between threat exposure and hippocampal and amygdala volumes in pediatric samples (6,27,28). Although neither hippocampal nor amygdala volumes were associated with depression, both moderated the association between stressful life events and depression, such that this relationship was significantly stronger among youths with smaller hippocampal and amygdala volumes. Further, we found evidence of moderated mediation, such that increases in depressive symptoms among youths exposed to violence were mediated by reductions in hippocampal and amygdala volumes, but only among youth who experienced more stressful life events. Reductions in hippocampal and amygdala volumes may therefore be important mechanisms underlying stress sensitization to depression.

Alterations in stress reactivity and emotion regulation may explain the role of hippocampal and amygdala volumes in stress sensitization. Smaller amygdala volume is associated with greater physiological reactivity to stress (6,7,30). This increased reactivity to acute, episodic stressors may contribute to greater vulnerability to depression over time. Adults with greater amygdala reactivity have been found to have a stronger association between stressful life events and depression symptoms 1-4 years later (31). Reductions in hippocampal volume may disrupt hippocampal regulation of the hypothalamic-pituitary-adrenal (HPA) axis, which mediates physiological responses to stress (8). Atypical HPA axis function is well-documented in individuals with depression (32,33). Reduced hippocampal and amygdala volume among youth exposed to violence ultimately contribute to heightened emotional reactivity in the face of novel stressors.

While this study has numerous strengths, some limitations constrain interpretability and suggest potential future directions for research. First, the distribution of the number and severity of stressful life experiences differed systematically between violence exposed and unexposed participants. While this is consistent with patterns in the general population, it leads to some differential restrictions to the range of stressful life events in the two groups. Next, because subcortical volumes were measured after violence exposure occurred, it is impossible to establish whether or not differences in hippocampal volume preceded the violence exposure. Prospective longitudinal designs among families at high risk for violence exposure could potentially more clearly establish the causal link between violence exposure and hippocampal and amygdala volumes. Indeed, one study has shown that an intervention aimed at enhancing supportive parenting in low socioeconomic status families with adolescent children prevented reductions of hippocampal and amygdala volumes (34). Finally, future work examining behavioral and neurophysiological indicators of hippocampal and amygdala functioning in relation to hippocampal and amygdala volumes and stress sensitization could further elucidate how reduced hippocampal and amygdala volumes acts as mechanisms of stress sensitization to depression and identify behavioral targets for intervention.

### Conclusion

Childhood violence exposure is associated with smaller hippocampal and amygdala volumes, which is, in turn, associated with increases in the emergence of depressive symptoms over time in youth who experience stressful life events. Smaller hippocampal and amygdala volumes are therefore a plausible mechanism of stress sensitization and have the potential to serve as a biomarker of risk following exposure to violence.

## Acknowledgments

This research was funded by the National Institute of Mental Health (R01-MH103291 to McLaughlin, F31-MH116559 to Lambert), an Early Career Research Fellowship from the Jacobs Foundation (McLaughlin), and a OneMind Institute Rising Star Award (McLaughlin). We are grateful to Debbie Bitran, Andrea Duys, and Azure Reid-Russell for help with participant recruitment and testing.

## References

1. McLaughlin KA, Conron KJ, Koenen KC, Gilman SE. Childhood adversity, adult stressful life events, and risk of past-year psychiatric disorder: A test of the stress sensitization hypothesis in a population-based sample of adults. Psychol Med. 2010;40(10):1647–58.

2. Kessler RC, McLaughlin KA, Green JG, Gruber MJ, Sampson NA, Zaslavsky AM, et al. Childhood adversities and adult psychopathology in the WHO World Mental Health Surveys. Br J Psychiatry. 2010;197(05):378–85.

3. Hammen C, Henry R, Daley SE. Depression and sensitization to stressors among young women as a function of childhood adversity. J Consult Clin Psychol. 2000;68(5):782–7.

4. Harkness KL, Bruce AE, Lumley MN. The role of childhood abuse and neglect in the sensitization to stressful life events in adolescent depression. J Abnorm Psychol. 2006.

5. McLaughlin KA, Weissman DG, Bitran D. Childhood Adversity and Neural Development: A Systematic Review. Annu Rev Dev Psychol. in press.

6. McLaughlin KA, Sheridan MA, Gold AL, Duys A, Lambert HK, Peverill M, et al. Maltreatment Exposure, Brain Structure and Fear Conditioning in Children and Adolescents. Neuropsychopharmacology. 2016;41(8):1956–64.

7. Yang RJ, Mozhui K, Karlsson R-M, Cameron HA, Williams RW, Holmes A. Variation in Mouse Basolateral Amygdala Volume is Associated With Differences in Stress Reactivity and Fear Learning. Neuropsychopharmacology. 2008;33(11):2595–604.

8. Frodl T, O’Keane V. How does the brain deal with cumulative stress? A review with focus on developmental stress, HPA axis function and hippocampal structure in humans. Neurobiology of Disease. 2013.

9. McLaughlin KA, Koenen KC, Bromet EJ, Karam EG, Liu H, Petukhova M, et al. Childhood adversities and post-traumatic stress disorder: Evidence for stress sensitisation in the World Mental Health Surveys. British Journal of Psychiatry. 2017.

10. Schmaal L, Veltman DJ, Van Erp TGM, Smann PG, Frodl T, Jahanshad N, et al. Subcortical brain alterations in major depressive disorder: Findings from the ENIGMA Major Depressive Disorder working group. Mol Psychiatry. 2016.

11. MacMillan S, Szeszko PR, Moore GJ, Madden R, Lorch E, Ivey J, et al. Increased Amygdala: Hippocampal Volume Ratios Associated with Severity of Anxiety in Pediatric Major Depression. J Child Adolesc Psychopharmacol. 2003.

12. Rosso IM, Cintron CM, Steingard RJ, Renshaw PF, Young AD, Yurgelun-Todd DA. Amygdala and hippocampus volumes in pediatric major depression. Biol Psychiatry. 2005.

13. Ellis R, Fernandes A, Simmons JG, Mundy L, Patton G, Allen NB, et al. Relationships between adrenarcheal hormones, hippocampal volumes and depressive symptoms in children. Psychoneuroendocrinology. 2019.

14. Caetano SC, Fonseca M, Hatch JP, Olvera RL, Nicoletti M, Hunter K, et al. Medial temporal lobe abnormalities in pediatric unipolar depression. Neurosci Lett. 2007.

15. Whittle S, Yap MBH, Sheeber L, Dudgeon P, Yücel M, Pantelis C, et al. Hippocampal volume and sensitivity to maternal aggressive behavior: A prospective study of adolescent depressive symptoms. Development and Psychopathology. 2011.

16. Gorka AX, Hanson JL, Radtke SR, Hariri AR. Reduced hippocampal and medial prefrontal gray matter mediate the association between reported childhood maltreatment and trait anxiety in adulthood and predict sensitivity to future life stress. Biol Mood Anxiety Disord. 2014.

17. Bifulco A, Brown GW, Harris TO. Childhood Experience of Care and Abuse (CECA): A Retrospective Interview Measure. J Child Psychol Psychiatry. 1994;35(8):1419–35.

18. Bernstein DP, Ahluvalia T, Pogge D, Handelsman L. Validity of the childhood trauma questionnaire in an adolescent psychiatric population. J Am Acad Child Adolesc Psychiatry. 1997;36(3):340–8.

19. Steinberg AM, Brymer MJ, Kim S, Briggs EC, Ippen CG, Ostrowski SA, et al. Psychometric Properties of the UCLA PTSD Reaction Index: Part I. J Trauma Stress. 2013.

20. Finkelhor D, Hamby SL, Ormrod R, Turner H. The Juvenile Victimization Questionnaire: Reliability, validity, and national norms. Child Abuse and Neglect. 2005.

21. Walker EA, Unutzer J, Rutter C, Gelfand A, Saunders K, VonKorff M, et al. Costs of health care use by women HMO members with a history of childhood abuse and neglect. Arch Gen Psychiatry. 1999.

22. Kovacs M. Children’s depression inventory (CDI2). North Tonawanda NY: Multi-Health Systems Inc.; 2011.

23. Hammen C. Self-cognitions, stressful events, and the prediction of depression in children of depressed mothers. J Abnorm Child Psychol. 1988;

24. Hammen C. Generation of stress in the course of unipolar depression. J Abnorm Psychol. 1991;100(4):555–61.

25. U.S. Census Bureau. Poverty Thresholds [Internet]. Available from: https://www.census.gov/data/tables/time-series/demo/income-poverty/historical-poverty-thresholds.html

26. Buuren S van, Groothuis-Oudshoorn K. mice□: Multivariate Imputation by Chained Equations in *R*. J Stat Softw. 2011;

27. Hanson JL, Nacewicz BM, Sutterer MJ, Cayo AA, Schaefer SM, Rudolph KD, et al. Behavioral problems after early life stress: Contributions of the hippocampus and amygdala. Biol Psychiatry. 2015;77(4):314–23.

28. Saxbe D, Khoddam H, Piero LD, Stoycos SA, Gimbel SI, Margolin G, et al. Community violence exposure in early adolescence: Longitudinal associations with hippocampal and amygdala volume and resting state connectivity. Dev Sci. 2018;21(6):e12686.

29. Espejo EP, Hammen CL, Connolly NP, Brennan PA, Najman JM, Bor W. Stress Sensitization and Adolescent Depressive Severity as a Function of Childhood Adversity: A Link to Anxiety Disorders. J Abnorm Child Psychol. 2007;35(2):287–99.

30. Trotman GP, Gianaros PJ, Zanten JJCSV van, Williams SE, Ginty AT. Increased stressor-evoked cardiovascular reactivity is associated with reduced amygdala and hippocampus volume. Psychophysiology. 2019;56(1):e13277.

31. Swartz JR, Knodt AR, Radtke SR, Hariri AR. A Neural Biomarker of Psychological Vulnerability to Future Life Stress. Neuron. 2015;85(3):505–11.

32. Burke HM, Davis MC, Otte C, Mohr DC. Depression and cortisol responses to psychological stress: A meta-analysis. Psychoneuroendocrinology. 2005;

33. Knorr U, Vinberg M, Kessing L V., Wetterslev J. Salivary cortisol in depressed patients versus control persons: A systematic review and meta-analysis. Psychoneuroendocrinology. 2010.

34. Brody GH, Gray JC, Yu T, Barton AW, Beach SRH, Galván A, et al. Protective Prevention Effects on the Association of Poverty With Brain Development. JAMA Pediatr. 2017 01;171(1):46–52.

35. McCabe CJ, Kim DS, King KM. Improving Present Practices in the Visual Display of Interactions. Adv Methods Pract Psychol Sci. 2018;

